# High Throughput Volume Flow Cytometry (parallel-iLIFE) Resolves Mitochondrial Network On the Go

**DOI:** 10.1101/2020.10.14.334466

**Authors:** Prashant Kumar, Prakash Joshi, Jigmi Basumatary, Partha Pratim Mondal

**Author notes:** Corresponding author (Partha P. Mondal).

## Abstract

Cell screening and viability studies are paramount to access cell morphology and intracellular molecular variations within large heterogeneous populations of cells. This forms the basis for diagnosis of infections, evaluating immunohistochemistry and routine histopathology. The proposed volume flow cytometry (also termed as, parallel Integrated Light-sheet imaging and flow-based enquiry (parallel-iLIFE)) is a powerful method that adds new capabilities (3D volume visualization, organelle-level resolution and multi-organelle screening) powered by light sheet based illumination. Unlike state-of-the-art point-illumination based imaging cytometry techniques, light sheet based parallel-iLIFE technique is capable of screening species with high throughput and near diffraction-limited resolution. The flow system was realized on a multichannel (Y-type) microfluidic chip that enables visualization of mitochondrial network of several cells in-parallel at a relatively high flow-rate of 2000 *nl*/*min*. The calibration of system requires study of point emitters (fluorescent beads) at physiologically relevant flow-rates (50−2000 *nl*/*min*) for determining flow-induced optical aberration in the system point spread function (PSF). Subsequently, recorded raw images and volumes were deconvolved with flow-variant PSF to reconstruct cellular mitochondrial network. High throughput investigation of HeLa cells were carried out at sub-cellular resolution in real-time and critical parameters (mitochondria count and size distribution, morphology and cell strain statistics) are determined on-the-go. These parameters determine the physiological state of cells and the changes in mitochondrial distribution over-time that may have consequences in disease diagnosis. The development of volume flow cytometry system (parallel-iLIFE) and its suitability to study sub-cellular components at high-throughput high-content capacity with organelle-level resolution may enable disease diagnosis on a single microfluidic chip.

## I. INTRODUCTION

Disease diagnosis and its cure requires single platform based diagnostic systems that are multifunctional and enables rapid analysis. Imaging flow cytometry is one such powerful system that is capable of addressing diverse study essential in fields ranging from cell biology to applied physics [1] [2] [3] [15]. These compact systems combine the strengths of imaging and flow cytometry. Apart from providing the basic functions of conventional cytometry to count, sort, mix and analyse cells, it also offers metrics that can be used to discern cells based on their morphology, cellular variations and organelle-level information [4] [5] [6] [7] [8] [9] [10] [11] [12] [13] [14]. This helps in specific and accurate decision making and has immediate implications in medicine, health-care and clinical diagnosis.

Conventional point-illumination based flow cytometers are constrained by low-throughput (due to sequential cell screening), poor resolution (owing to flow-induced optical aberration and limited photon budget) and complex instrumentation (due to the necessity of sheath fluid based hydrodynamic focussing). A light sheet based technique is developed that is capable of overcoming these limitations. Unlike point based flow cytometry that requires hydrodynamic focussing of specimen to a narrow space in order for it to be intersected by the system PSF, the sheet illumination sections the entire flow channel. This brings the advantage of large cross-section interrogation of the specimens in-parallel (upto 1000 cells per minute) during flow [16] [17]. Light sheet imaging requires a widefield based orthogonal detection that can resolve features with diffraction-limited resolution [15] [18] [19] [20]. The geometry of light sheet illumination eliminates the complexity associated with hydrodynamic imaging. Moreover, the technique facilitates high quality volume reconstruction of the entire specimen during flow. The sectional images of the specimen are deconvolved and stacked together in real-time to reconstruct the volume-stack, a feature never reported before. So, parallel iLIFE system builds on sheet interrogation to map the sectional 2D images of the target specimen on to the 3D space for volume reconstruction. Since the first demonstration on simple microfluidic chip (straight micro-channels) in 2013 [16] [17], it has brought in paradigm-shift from point to plane based interrogation. Subsequently, the system was modified and advanced to image fluorescently-labelled organelles in live worms (C. elegans) [15]. In the past few years, lightsheet imaging cytometry (LIC) has gained popularity and many research groups have started exploring LIC to incorporate live organism imaging [21], time-stretch imaging [22], phase-contrast [23], high-speed interrogation [24], fast sectioning [25], depth-penetration [26], determination of protein concentration [27], measurement of intracellular content [11] and label-free cell identification [28]. The new volume flow cytometry (parallel-iLIFE) is the next generation technology for applications in diverse fields.

## II. RESULTS

The development of multichannel volume flow cytometry is advantageous when compared to traditional imaging cytometry systems. This helps in determining critical parameters that may pin-point the underlying biological mechanism, understand disease progression and may assist in drug therapy.

### A. Light sheet based parallel iLIFE System

Existing imaging cytometry system requires much more than the existing functionalities (count, sort, mix and filter) for it to become a full-fledged diagnostic health-care system. Towards this goal, the parallel iLIFE system is developed to add more functionalities such as, high resolution cross-sectional imaging, real-time volume reconstruction, biophysical parameter estimation and parallel screening of multiple specimens. This will further lead to the understanding of cell morphology, intracellular protein concentration, organelle distribution and their changes from normal cell type indicating diseased state.

The parallel-iLIFE cytometry system consists of three major sub-systems: (1) Light sheet based illumination sub-system, (2) Microfluidic chip based cell flow platform and, (3) Orthogonal widefield detection sub-system. The illumination system is the heart of iLIFE system that uses a sheet-of-light to illuminate the specimens flowing through the sample platform. Unlike the existing state-of-the-art cytometry systems, light sheet illumination is significantly different that employs a sheet-of-light rather than point-PSF. This is a paradigm-shift that lifts a lot of constraints (such as, the requirement of hydrodynamic focussing and sequential specimen interrogation) and related complications faced by existing cytometers. This brings us to the second important aspect of specimen flow system. The existing cytometers require sheath fluid that hydrodynamically focus and cause the cells to align in the direction of flow. The proposed parallel iLIFE system employs a microfluidic chip based cell flow that does not require cell alignment. The sheet is chosen large enough so that it can cross-section the entire micro-channel array. The detection is carried out at 90° in a widefield mode which gives the system an unprecedented near diffraction-limited resolution.

Figure 1 shows the schematic diagram of actual parallel-iLIFE system. The system is used in a fluorescence mode for the proposed study related to organelle (mitochondria) imaging. Laser light of wavelength 532 *nm* with a beam-width of 1.5 *mm* is expanded using a standard beam expander to expand the beam 5-times. This is essential to fill the back-aperture of cylindrical lens (CL) to enable use of maximum available aperture angle. The resultant beam is 1D focussed and allowed to fall at the back-aperture of objective lens (*O*_*ill*_). This gives rise to diffraction-limited light-sheet PSF at the focus of objective lens. The sheet thus generated intersects the near-transparent PDMS based microfluidic chips containing an array of Y-type micro-channels. The cells are stored in the reservoir (R) which is connected to one end of channel array and the flow is generated by flow-pump operating in a withdrawl-mode. This results in a continuous flow of cells at a controlled flow. The mitochondria of cell was labelled with Mitotracker orange (*Ex*: *Em* = 554 *nm*: 576 *nm*) dye. This enabled us to observe Mitochondrial network in the 3D cell. The emission of fluorescence from cells flowing through the array of micro-channels is collected by the detection objective placed at 90°. The light is subsequently focussed by the tube lens (TL, *f* = 125 *mm*) to the fast CCD camera (maximum frame rate of 2.3k frames / sec). On its way to detector, the light is filtered by a combination of filters (bandpass and notch filters) placed in the filter box (F-Box). The inset of figure 1 shows a schematic of intersecting cell flow system with light sheet.

**FIG. 1:**
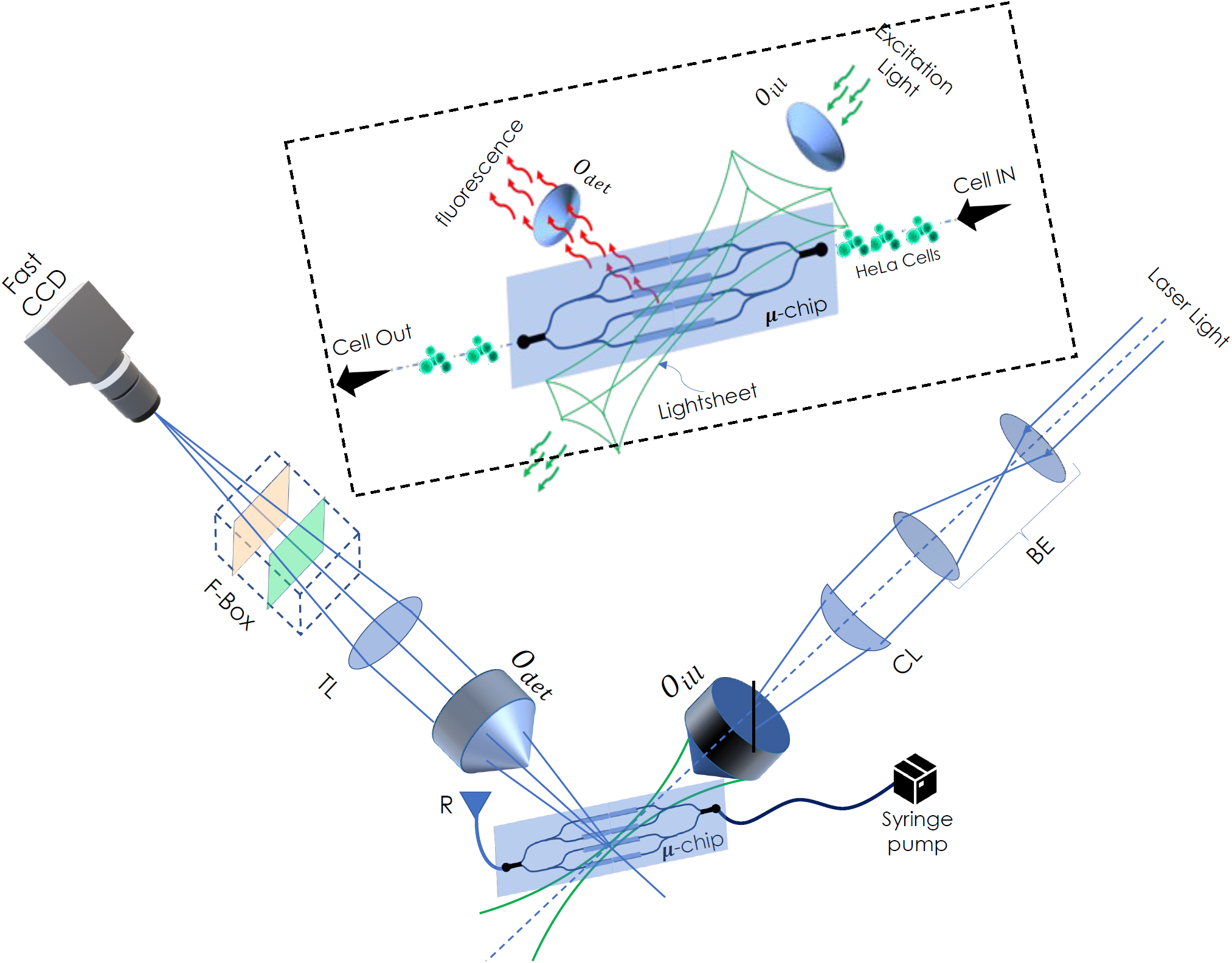
Schematic diagram of a volume flow cytometry system (termed as, parallel-iLIFE). Both illumination and detection subsystems are configured in an orthogonal setting for light sheet based interrogation. A laser of 532 *nm* light is expanded (using beam-expander, BE) and subjected to cylindrical lens (*f* = 150 *mm*) - objective lens (10X, 0.25 NA) combination. This generates diffraction-limited light sheet that sections the entire Y-type 4-microfluidic channel array. Several cells flowing through the channels gets simultaneously interrogated by light sheet. The fluorescence from the labelled mitochondria is collected by detection objective (20X, 0.4 NA), filtered by filter-box (containing a set of longpass and notch filters) and focussed to the CCD camera by tube-lens (TL). Instant deconvolution is carried out and cell volume is reconstructed. The inset shows an expanded view of light sheet intersecting the microfluidic channel-array.

### B. Dynamic Light Sheet Characterization

The light field characterization and its interaction with the specimens flowing through PDMS are at the heart of illumination sub-system. This inherently determines the selectivity and FOV of the imaging system. A focused light sheet reduce the plane thickness thereby improving cell sectioning ability and boosts signal-to-background ratio. Fig. 2 shows experimentally measured light field at and around the focus of illumination objective (*O*_*ill*_). The field is measured directly by translating a camera (equipped with ND filters) in the beam-path. A CCD camera was placed in the beam-path and z-scanned along the optical axis to record the field at each *z*-increment (see, Fig.2). We have employed step-gradient neutral density (ND) filters to avoid saturation of intensity field at the camera. The scanning was performed by a translator carrying the camera. The 3D recorded field along with its XY, YZ and XZ views are shown in Fig.2. In addition, field characterization is carried out using intensity plots along X Y and Z axes to determine its size and shape. The shape of the light sheet can be assessed from the associated 2D (XY, YZ and XZ) images. Intensity plots show that the generated light sheet has a dimension of ≈ 480 × 176 *µm*^2^ along (*y* × *z*) (full-width at 25% maximum) and thickness, 3.75 *µm*. 3D view clearly display the planar light sheet field at and around the geometrical focus. Specifically, strong confinement of the field is evident along the z-axis resembling sheet-like field structure. The intersection of light sheet field with the flowing cells determines the reconstruction quality of cell volume.

**FIG. 2:**
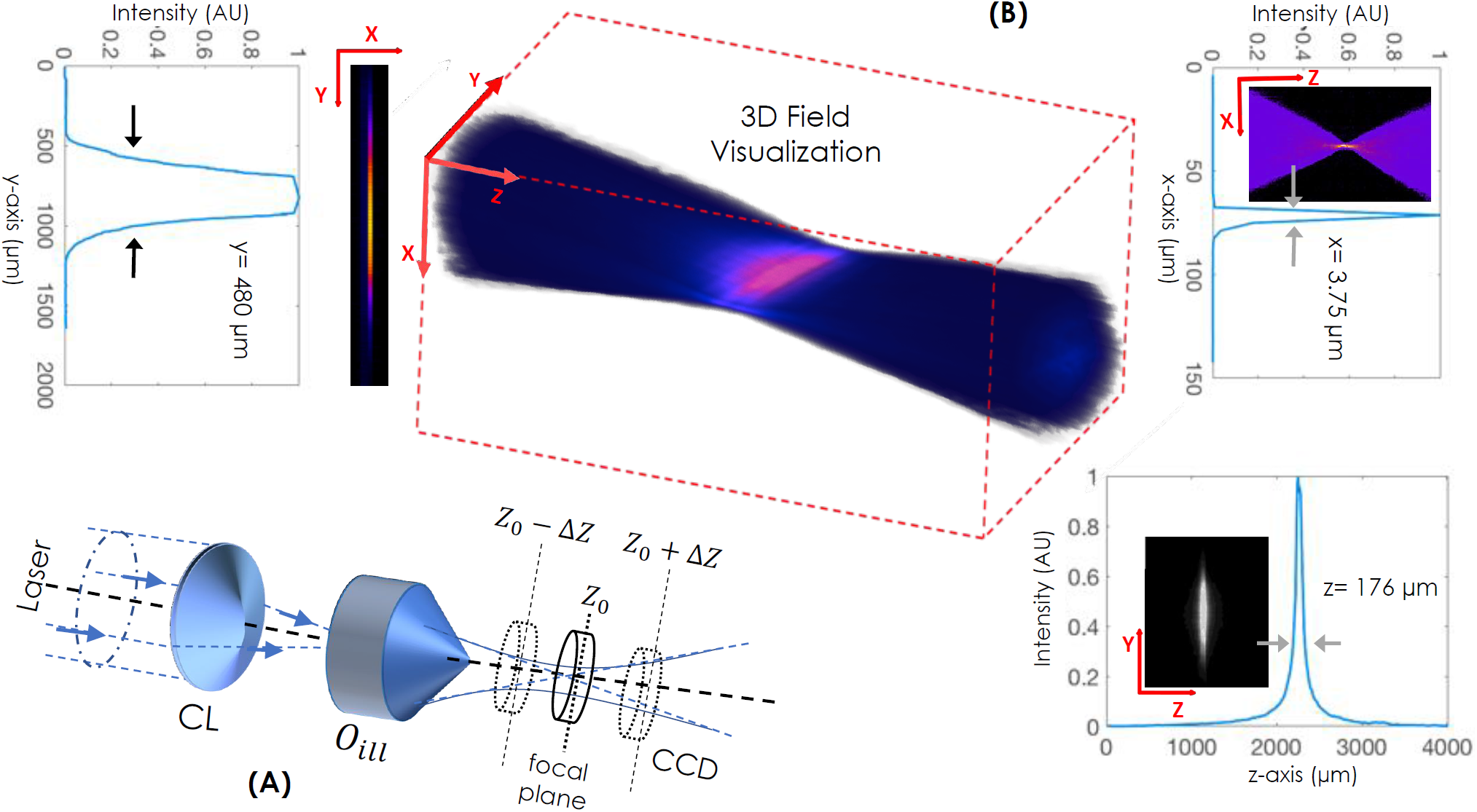
3D Field Visualization: (A) Optical set-up to measure the diffraction-limited light sheet field. The CCD camera is placed on a linear-translator and scanned along the optical z-axis to record 2D field intensity. (B) The field images are stacked together to reconstruct 3D field. XY, XZ and YZ views of three dimensional light sheet field are also shown. The corresponding dimensions of experimentally recorded light sheet shows a light sheet of dimension ≈ 480 × 176 *µm*^2^ along (*y* × *z*) and has a thickness of 3.75 *µm*.

### C. Flow based Dynamic PSF

The parallel-iLIFE is a dynamic flow imaging system that requires accurate characterization of system PSF. This has profound implication on the image quality. The very fact that a moving target induces flow-based distortion compromises system resolution and thereby affects overall quality of sectional images and cell volume. To determine system PSF during flow, we have used fluorescent beads (of size, ∼ 1 *µm*) as point source flowing through the channel at different flow-rates. Figure 3 shows fluorescent beads (*Ex*: *Em* = 535 *nm*: 575 *nm*) at flow-rates ranging from 50 *nl*/*min* to 2000 *nl*/*min*. It may be noted that, parallel-iLIFE is operated in the mentioned dynamic flow range for the present study. At low flow-rates, the system PSF is unperturbed and shows negligible motion-blur due to flow. The situation however changes as the flow-rate is increased to 1000 *nl*/*min*, but it becomes visually apparent at around 2000 *nl*/*min*. To characterize the effect of flow, the intensity plots across the cross-section of bead is plotted for the entire dynamic range. Subsequently, quantification is carried out by fitting Gaussian and the apparent size (due to motion-blur) of system PSF size is determined along flow axis. The beads flowing through the channel are found to be elongated along the flow direction, with its apparent size increased by almost ≈ (*σ*_2*k nl*/*min*_/*σ*_50 *nl*/*min*_) = 1.8645 fold. This is quite significant considering the fact that we are seeking high resolution. Throughout the experiment, we have used a frame-rate of 38 frames / sec to collect more photons. Large frame-rates reduce both motion-blur and system resolution (as determined by the number of detected photons). So a balance between flow-rate and camera frames / sec is necessary for quality volume reconstruction. The linear relationship between the PSF size and flow-rate is quite evident from the straight line fit. A small slope of 1.39 × 10^−3^ *µm*/*nl*/*min* ensures that the system operates in linear regime. All these lead us to believe that, flow in the considered imaging regime (50 *nl*/*min* to 2000 *nl*/*min*) is linear in nature and lead to proportional changes in the size of system PSF along the flow direction.

**FIG. 3:**
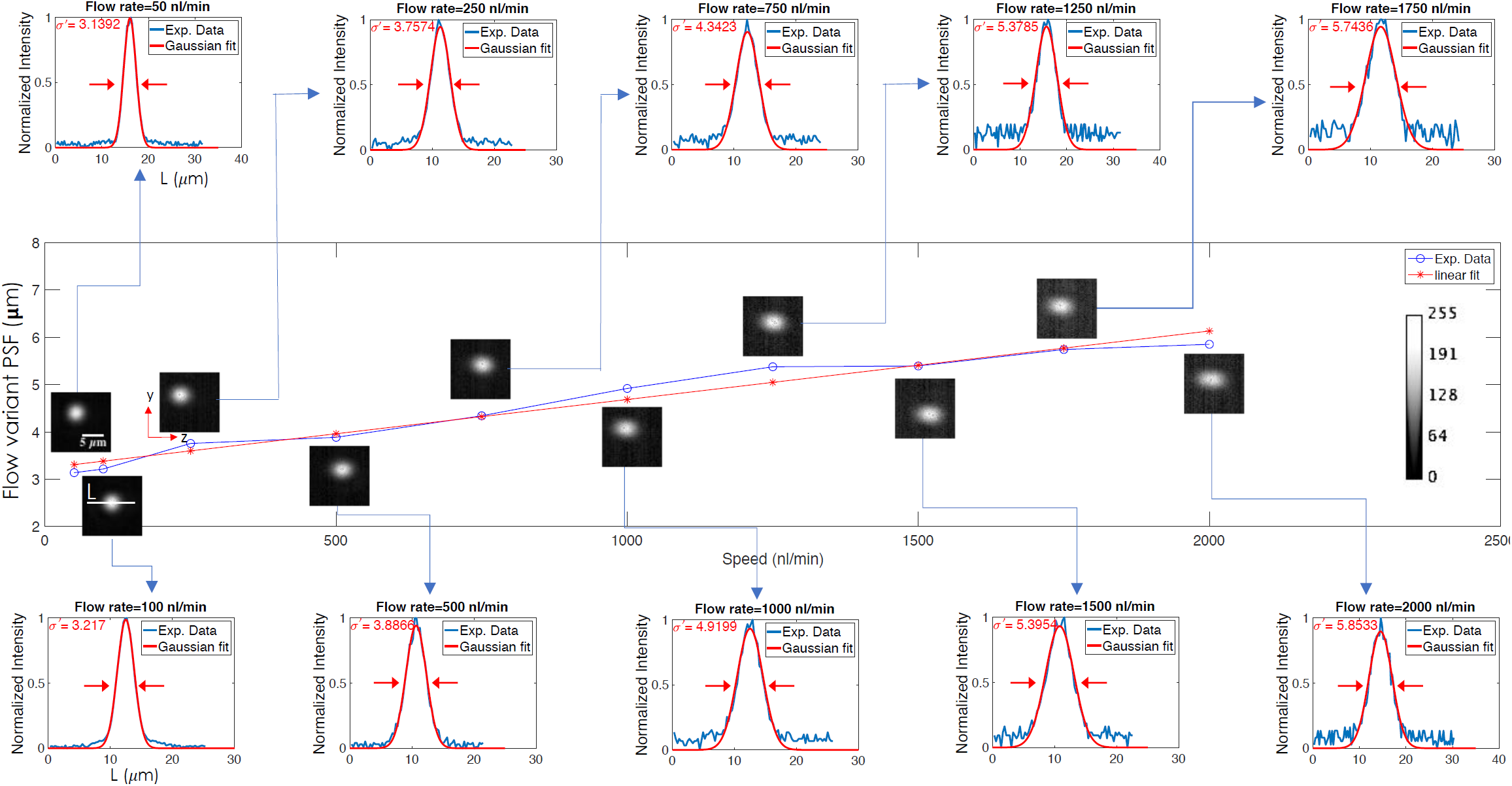
Flow-variant System PSF: The PSF at different flow rates ranging from 50 *nl*/*min* to 2000 *nl*/*min* is shown. Fluorescent beads of size 1 *µm* is used as point source for determining PSF. With increased flow, the point sources appeared elongated along the flow direction for which the cross-section (across the PSF indicated by white line *L*) is plotted (see, blue curve). To evaluate the apparent elongation caused by motion-blur, the data is fitted with a Gaussian function (see, red line) and the corresponding full-width at half-maximum (FWHM) is calculated. Subsequently, flow-variant PSF size 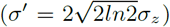 is plotted at different flow rates that indicate a linear increase in apparent size of system PSF.

### D. Simultaneous Multichannel Imaging of Beads and HeLa Cells

Simultaneous real-time volume imaging of specimens flowing through multiple microfluidic channels is a daunting task. This paves the way for next-generation flow based diagnostics. The parallel iLIFE system is designed to address this issue and overcome the constraints associated with existing state-of-the-art imaging cytometry systems. The key features that distinguish parallel-iLIFE with the existing imaging cytometry system are: (1) multichannel-array based imaging, (2) large cross-sectioning ability, (3) high throughput interrogation (due to microchannel array assembly) and volume rendering in real-time. The choice of cylindrical lens - objective lens combination is critical in determining the size of light sheet. We have chosen this combination so that the light sheet cross-sections the entire microchannel array, sectioning the flowing specimens simultaneously in all the channels. This ensures parallel interrogation of cells flowing through it, giving it the much needed advantage of high throughput interrogation along with real-time single cell volume reconstruction. Figure 4 show fluorescent beads and HeLa cells (fluorescently-labelled mitochondria) flowing through single, double and four channels. As a first step, we have flown fluorescent beads to determine dynamic flow parameters such as, flow-rate and optimal sample concentration among others. We observed that the channels often get blocked at low flow-rates (< 600 *nl*/*min*) predominantly due to gravity. Repeated study shows that a high flow rate of 2000 *nl*/*min* is appropriate for smooth cell-flow, owing to its large mass (compared to fluorescent beads) and relatively large size (diameter ∼ 15 − 25 *µm*). Cell samples were freshly prepared and loaded to the sample reservoir (R in Fig.1) and the suction-pump is operated at 2000 *nl*/*min*. Details of frame-rate, exposure time and other critical details are discussed in the Methods section. Few cross-sectional images along with the associated intensity plots are also shown in figure 4. The plots show internal distribution of mitochondria in a flowing cell along chosen lines (see, white line). The discrete presence of mitochondria / mitochondrial assembly (see, arrows in the intensity plot) determines the practical resolution of our flow cytometry system. This suggests that parallel-iLIFE is capable of imaging mitochondria distribution in a large population of cells.

**FIG. 4:**
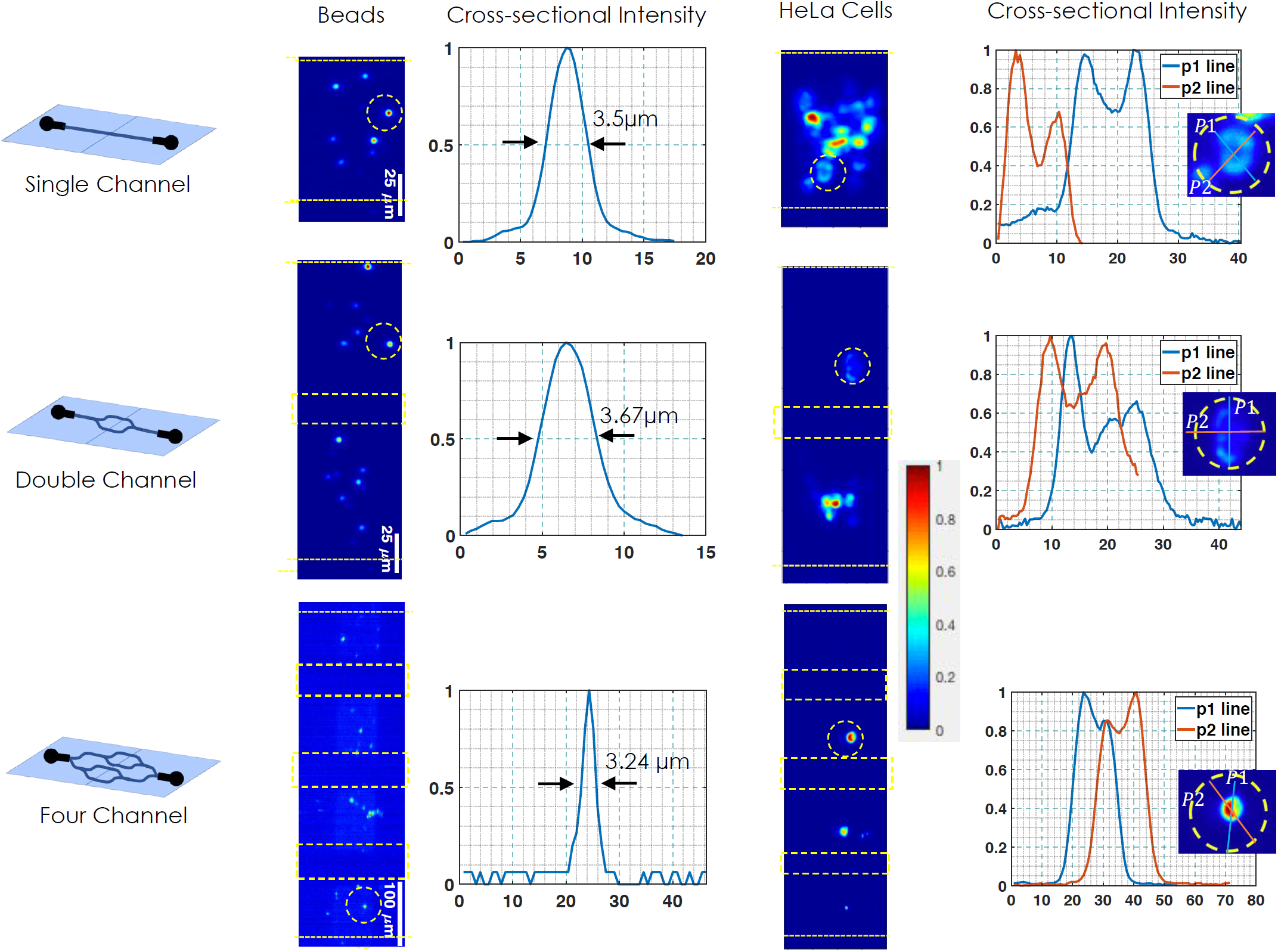
Parallel Multichannel Visualization: (A) Fluorescent beads flowing through different channel types (1, 2 and 4 channels). Beads flowing through the channels are simultaneously illuminated by light sheet and imaged by fast CCD camera. Corresponding intensity plots show FWHM values of beads flowing through the channels. (B) Mitochondria-labelled HeLa cells flowing through different channel types and simultaneously imaged in-parallel as they pass through the light sheet, producing 4-8 cross-sectional images. The associated intensity plots show variations indicating mitochondrial network inside the cell.

### E. Count Statistics

Population study of a cell ensemble is critical to disease diagnosis and helpful to periodically access its progression. The proposed parallel-iLIFE has the potential for high throughput imaging. The microfluidic array geometry enables multifold parallel interrogation of diverse cell population. Figure 5 shows the count statistics for both the fluorescent beads and HeLa cells at varying flow rates. For beads, we observed a healthy double-fold increment in count statistics for double channel array while the increment is about 2.5 for 4-channel array. Incredibly, this is true for a dynamic range of flow-rates (from 50 *nl*/*min* to 2000 *nl*/*min*). However, the statistics for larger specimens such as HeLa cells, the count is found to be appreciable at low flow-rates (∼ 200 per minute at 500 *nl*/*min*) and better at large flow-rates (> 800 per minute at 2000 *nl*/*min*). However, variations are observed at intermediate flow-rates. This can be attributed to cell-accumulation near the junctions and gravity enforced settling of cells that restricts its mobility (due to large mass and size). Nonetheless, we noted a healthy cell count of > 800 for 4-channel array at large flow-rates. Additional constraint associated with HeLa cells are associated with local channel constriction and the affinity of HeLa cells to form clusters. These processes are found to be less prominent at large flow-rates. This is primarily the reason behind relatively large count at high flow-rates (2000 *nl*/*min*) when compared to relatively low flow-rates (say, 500 *nl*/*min*). Our results favor imaging cytometry at large flow-rates for cell cytology.

**FIG. 5:**
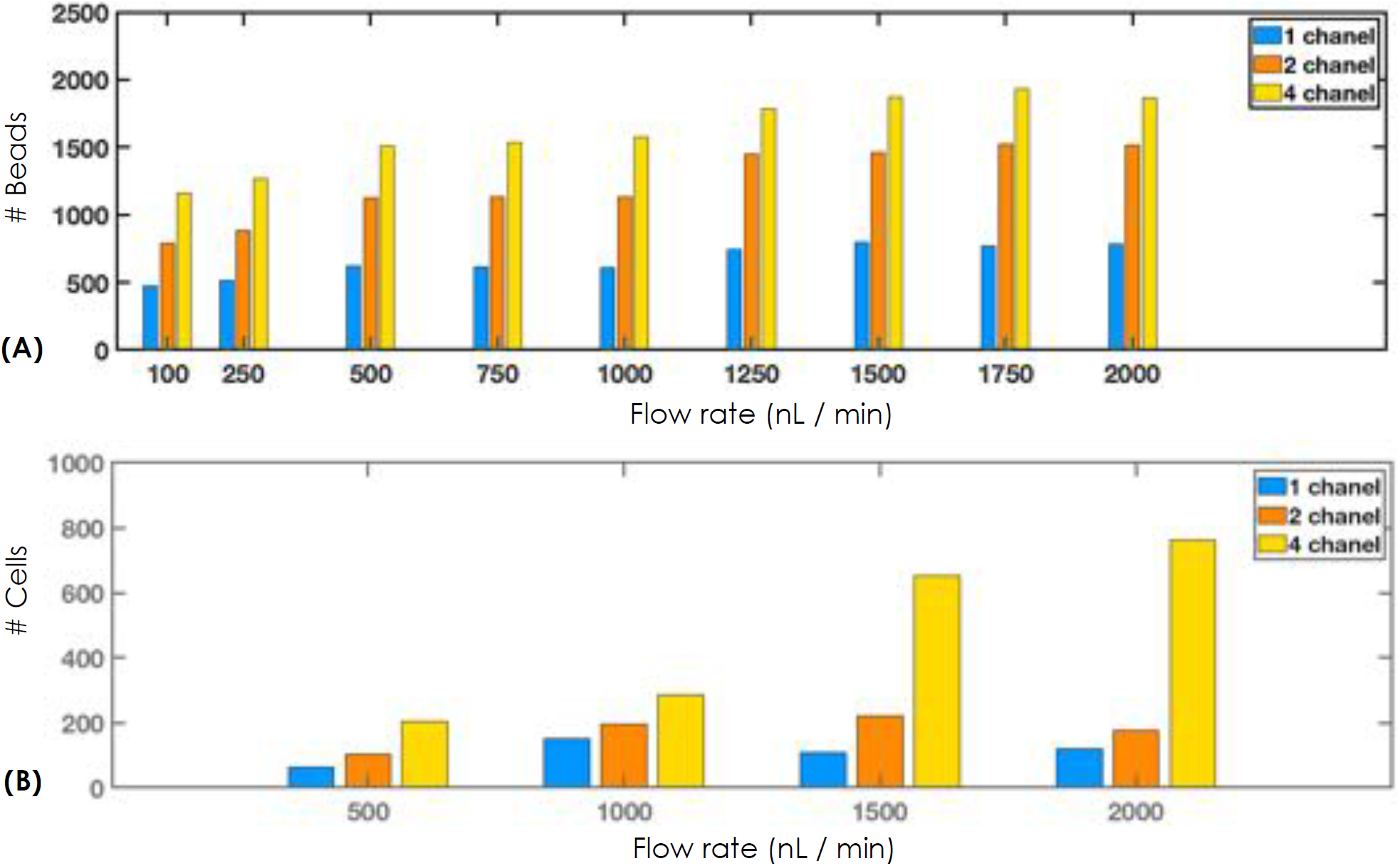
Counting Statistics for Volume Flow Cytometry System: (A) Fluorescent bead counting for 1, 2 and 4 channels showing a linear behaviour with increasing flow-rates, (B) HeLa cell counting shows a rather non-linear behaviour for 1 and 2 channels but displays a linear nature for 4-channels. This can be attributed to cell-clustering.

### F. Flow based Visualization of Mitocondrial Distribution in HeLa cells

Visualization of intracellular organelle at high speed and discerning its distribution is a step forward in imaging flow cytometry. The cells interrogated by the sheet illumination gives cross-sectional images of HeLa cells as they pass through the light sheet. Figure 6 shows the process of obtaining sectional images of multiple cells at a time and simultaneously in all the 4-channels. These 2D images are stacked together to obtain the entire 3D cell. Subsequently, the volume-stack is subjected to image reconstruction process (Deconvolution and background subtraction) to obtain 3D distribution of mitochondria in a single HeLa cell. Schematically the process is shown in figure 6 along with the distribution of mitochondrial network for few HeLa cells (see, Fig. 6B) along with its morphology (see, Fig. 6C). The distribution of mitocondria and its morphology in cell are key parameters for determining diseased state and rapid growth of HeLa cells.

**FIG. 6:**
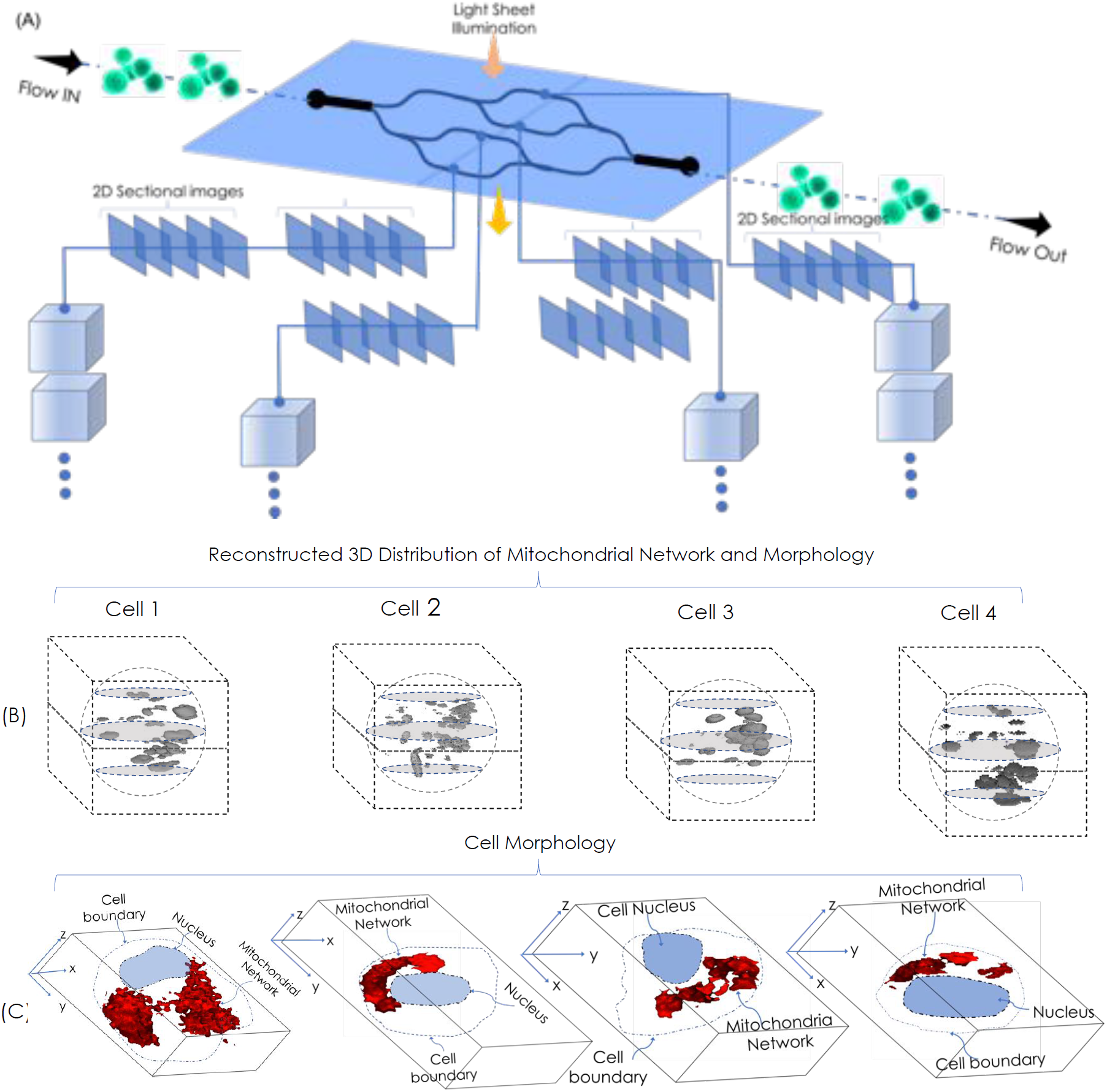
Parallel ilIFE Imaging and Volume Reconstruction: (A) A cartoon of iLIFE data acquisition system beginning from cell interrogation to volume reconstruction on the go. (B) 3D reconstructed cells displaying mitochondria distribution inside flowing Hela cells. (C) Reconstructed volume showing mitochondrial network (displayed in red color) inside HeLa cells. For clarity, nucleus and cell boundary are drawn. The reconstruction show organelle-level resolution in flow imaging system.

### G. Biophysical Parameter Estimation

Mitochondria continuously divide and fuse and their size and distribution are finely tuned [30] [31]. The fission and fusion of mitochondria play key roles in maintaining its integrity. Moreover, distribution of mitochondria contributes to the functioning of organelles that lead to healthy cell growth and its survival. Diseases such as cancer and Alzheimer are known to disturb this balance, causing change in size distribution and their relative count. Parallel iLIFE system has the ability to extract these parameters and related statistics for a large population of diseased cells leading to organelle-level investigation. Such a system is capable of assisting disease diagnosis both at sub-cellular and inter-cellular level. To demonstrate the capability of parallel iLIFE system, three important biophysical parameters (local mitochondria density, size distribution and cell strain statistics) were determined. Figure 7 shows estimated parameters at varying flow-rates. Fig. 7A shows the actual count of mitochondria / clustered-mitochondria per cell. It is evident that the count is high at low flow-rates and decrease monotonically at large flow. This is predictable due to the fact that large flow-rate results in motion-blur and high background that eventually makes it hard to detect small-to-medium sized mitochondria. But it is useful to access the relative count at specific flow-rates. Fig.7B shows the corresponding area distribution indicates mitochondria area peaking at 15 *µm*^2^. It is worthwhile to note that area HeLa cell lies somewhere between 300 − 700 *µm*^2^. In addition, cell strain studies are carried out that indicate the elasticity of cell and its deformability at large flow-rates. Post-detachment from the surface, cells assume the shape of a sphere. During flow it elongates along flow direction and consequently shrinks along perpendicular direction, assuming the shape of an ellipsoid with *a* and *b* as major and minor axis respectively. Thus, the metric *S*_*σ*_ ∝ (*a*/*b*) is taken as the measure of strain / deformation. Fig. 7C shows cell strain from low (500 *nl*/*min*) to high (2000 *nl*/*min*) flow-rates. A linear increase in strain is noted. Large deformation along flow direction helps in recording more optical sections during high flow (∼ 2000 *nl*/*min*) than at low flow (500 *nl*/*min*). Associated cell section statistics show that the camera is able to record large number of sections. The proportion of such cells have jumped to 52% at 2000 *nl*/*min* from 21% at 500 *nl*/*min* indicating large strain in a large population of cells. To our knowledge, multi-parameter estimation is unknown in imaging flow cytometry, and may pave way for improved health-care and diagnosis.

**FIG. 7:**
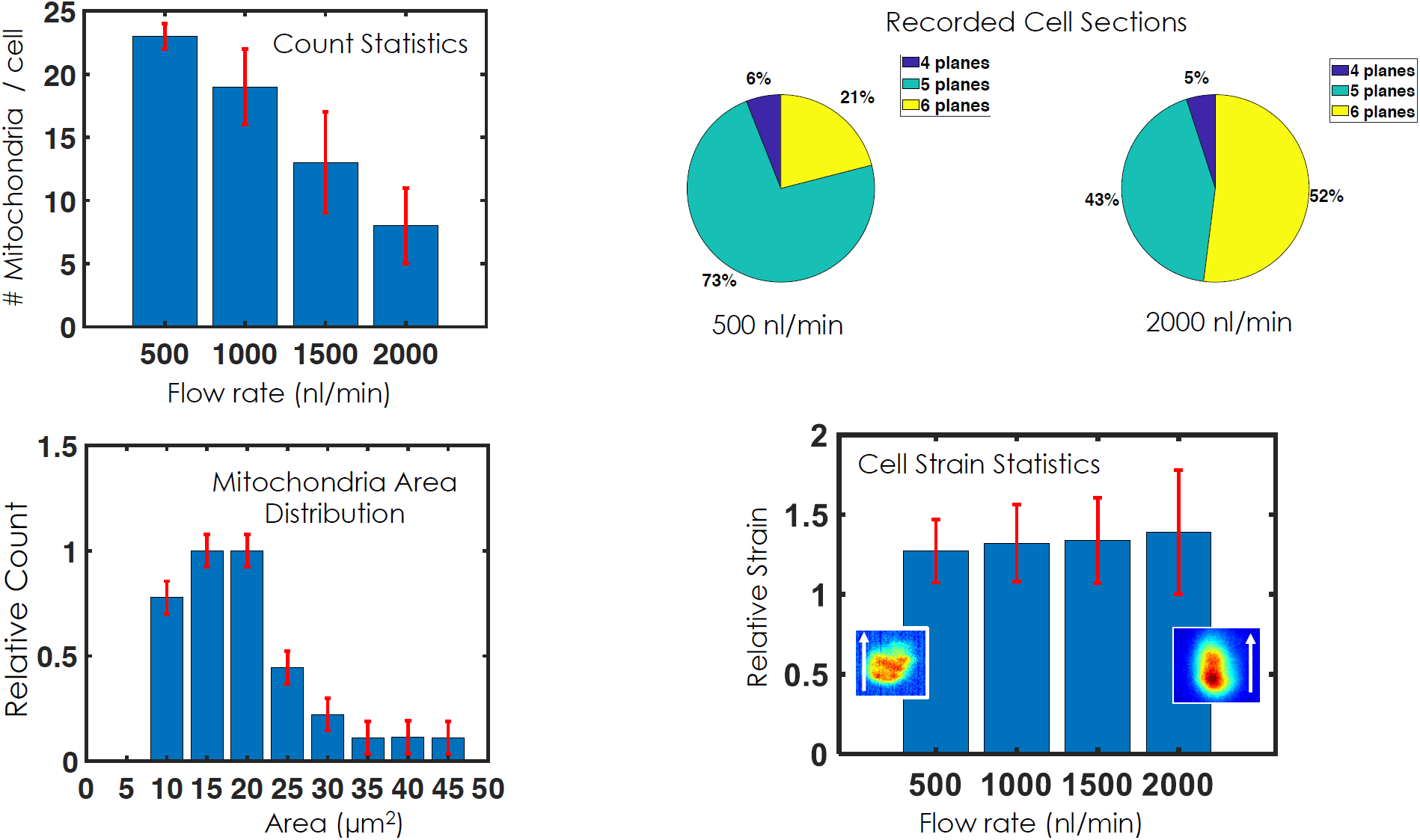
Biophysical Parameter Estimation: Mitochondria count statistics, size / area distribution and relative cell stress statistics from low (500 *nl*/*min*) to high (2000 *nl*/*min*) flow-rates. Count statistics show decrease in mitochondria count with increase in flow-rate. Area statistics show peak at 15 *µm*^2^. Strain statistics show linear increase in strain with flow-rate. A typical image of cell is also inserted along with the flow direction (indicated by white arrow) for visualization purpose only. Cell statistics is also displayed that show increase in the number of cross-sectional planes detected per cell at high flow-rates.

## III. DISCUSSION

The ability to obtain high resolution volume stacks of a large population of cells with high throughput has attractive applications in a variety of research fields, health-care and industry. Data obtained by parallel-iLIFE imaging cytometry contain cellular, multicellular and intracellular information of a large population, which can be used to classify cells types and access its state (disease / healthy). Moreover, the data can also be used for multiparameter analysis and gather large population statistics. Parallel-iLIFE cytometry may enable cancer cell identification [32][29], drug screening [33] and organelle distribution in cellular systems [34].

The primary advantage of parallel-iLIFE cytometry is its ability to reconstruct volume at high throughput (> 800 *cells*/*min*) with sectional-selectivity and near-diffraction-limited spatial resolution. The ability to generate numerous single-cell images in a short period of time has both advantages and disadvantages. This necessitates a delicate balance between image quality (resolution) and flow-speed of the specimen. The fact that high speed introduces motion-blur that compromises image resolution and low speed undermines high-throughput interrogation necessitates a balance between high throughput and spatial resolution. Other consequence of low speed is gravity-induced cell accumulation at the bottom of microfluidic channel and tendency of cells to form clusters.

In comparison to the existing state-of-the-art imaging cytometry techniques, parallel-iLIFE enables on-the-go volume visualization of organelles in single cell, high throughput interrogation (at approximately, 800 *cells*/*min*) and distribution of cellular organelles (mitochondria) in cells. Currently, parallel-iLIFE is the only volume flow cytometry method that has a cell throughput comparable to that of existing flow cytometry, indicating its suitability for analysing large population of cells for both intra- and inter-cellular investigations. Multiple channels facilitate interrogation of large cell population in parallel at moderate flow rates without effecting high throughput and thus prohibits degradation of system resolution caused by motion-induced blur. Here, parallel iLIFE is used to investigate mitochondrial distribution of a large population of cancerous HeLa cells. The fact that organelles can be fluorescently-labelled adds value and expands its horizon beyond conventional flow cytometry. Some of the exciting fluorescence based studies include, fluorescence imaging by frequency division multiplexing [35] [36] and time-delay integration [37] [38], where fluorescence images have played key role in determining critical parameters. Low-cost implementation of this new technology may extend its use in other fields as well.

The conventional imaging flow cytometry system use point focussing optics to interrogate the flowing specimens intersecting the point-like PSF. This requires complex instrumentation for hydrodynamic flow-focusing making it bulky and cumbersome. As a result, the system is capable of interrogating cells one-at-a-time in a sequential manner. Such an arrangement leads to low throughput interrogation and selective information gathering (related to cell shape and morphology) takes a back seat. In contrast, parallel-iLIFE cytometry uses a sheet of light to illuminate the specimen and a multichannel microfluidic chip for controlled flow of cells. This eliminates the need for hydrodynamic focussing and promotes simultaneous interrogation of multiple cells at a specific point in time. This is a huge advantage considering the slow nature of sequential scanning used in conventional flow cytometry system. Specifically, parallel-iLIFE system goes on to show accelerated imaging flow cytometry by simultaneously imaging specimens in multiple channels (Y-type channel array). In general, volume imaging is not possible with the existing cytometry system due to point-PSF. However, proposed volume cytometry system is inherently designed to take cross-sectional images primarily due to sheet-illumination and enables real-time volume reconstruction. It may be noted that, convolution of individual 2D sectional images is involved before they can be stacked together to obtain volume (see, Methods section). This requires the knowledge of system PSF. Point sources (fluorescent beads) are flown at varying flow-rates to determine the PSF-spread due to motion-blur. The PSF-distortion is found to vary linearly with flow in the range 50 *nl*/*min*. − 2000 *nl*/*min*. (see, Fig.3) limiting the number of recorded images obtained per cell somewhere between 4 − 8. Thereby flow-speed limits the information gathering capacity of parallel-iLIFE system. The resultant volume cytometry system is simple, miniaturizable and easily integrable with computational methods for high quality analysis (sub-cellular parameter estimation).

Our findings further suggest intracellular resolving capability of the proposed volume cytometry system. An orthogonal widefield detection predicts near diffraction-limited resolution, but we seldom reached this limit. This is primarily due to the optical aberration (motion-blur) caused by flow in the microfluidic channels. Consequently, the resolution of practical system is scaled and influenced by flow. Our results show that the factor is somewhere between (1.0 − 1.86), with the larger value for 2000 *nl*/*min*. This factor introduced by flow lowers the resolution of system by a factor of ≈ *η*_*f*_ = 1.86 of that of true diffraction-limited resolution of (λ/2*NA*) × *η*_*f*_ ≈ 1.979*µm*. Considering flow-based system, the resolution is quite good to investigate intercellular organelle of the size of few microns. Knowing that the size of mitochondria in a cell vary somewhere between 0.5 − 5*µm* (see, Fig. 7), we can safely estimate the resolution of our system to be in sub-micron to few micron range. This studies show that flow-based parallel iLIFE systems has good resolution for organelle-level investigation on the go.

The lightsheet illumination coupled with high-throughput flow system (Y-type multichannel microfluidic chip) and orthogonal widefield detection has enabled extraction of biophysical parameters (cell volume, mitochondrial network visualization and its distribution, local size distribution and cell strain) in flowing HeLa cancerous cells for further analysis (see, Fig. 7). The access to organelle-level resolution coupled with a healthy count of 800 *cells*/*min* (see, Fig.5) and multi-parameter estimation makes parallel-iLIFE a unique tool that combines the benefits of high-resolution microscopy, volume visualization and high-throughput cytometry.

## IV. METHODS

### A. Optical Setup

Parallel iLIFE imaging cytometry system has three major sub-systems: lightsheet illumination, multichannel microfluidic chip (specimen holder) and high-speed detection:

#### Lightsheet Illumination

Simultaneous illumination of multiple microfluidic channels allows interrogation of a large population of cells. A large light sheet is generated in order to cross-section the entire array of micro-channels through which cells were flown. A laser (532 nm Excel Laser, Quantum Lasers, UK) of wavelength 532 *nm* and beam-width 1.5 *mm* is used as the light source. The beam is expanded using a beam-expander (consists of two biconvex lens of focal-lengths 25 *mm* and 125 *mm* procured from Thorlans, USA) by a factor 5X-times. This is essential to just over-fill the back-aperture of cylindrical lens (*f* = 150 *mm*, Edmund Optics, Singapore) in order to utilize its full NA. The expanded beam is the 1D-focused by the cylindrical lens to form a horizontal light-sheet. An objective lens (Olympus 10X, 0.25 NA) is placed after the cylindrical lens to generate diffraction-limited vertical light sheet. The light sheet thus generated serves as the illumination PSF of parallel-iLIFE system.

#### Microfluidic Chip based Sample Flow System

Cells were flown through the Y-shapped microfluidic channel array as shown in Fig. 1. The chip is fixed to a home-bult chip holder and placed on a XYZ-translator for precise position with respect to light sheet for optically sectioning the entire channel array (an array of 4 channels of size 100*µm*^2^). The channel inlet is linked to sample reservoir containing the cells whereas, the outlet is connected to a flow-pump. The pump is operated in a suction-mode and the operations are controlled by computer based interface software. Both fluorescent beads (Fluospheres, Invitrogen) and HeLa cells (labelled with Mitotracker) were flown through the channel-array and imaged in-parallel.

#### Fast Detection

The mitochondria in the cell was labelled using mitotracker orange dye (Invitrogen) that has an emission maximum at 576 *nm*. The fluorescence from specimen (HeLa cells) is collected by the detection objective (Meiji, 20X, 0.4 *NA*). Subsequently, the light is filtered by a set of filters (notch filter (ZET532nf purchased from Chroma) to remove the illumination 532 *nm* light and a long-pass filter (purchased from Thorlabs) to filter-out the background to retain the fluorescence from cell mitochondria. The filtered light is then focused to the camera chip (pixel size ∼ 5.5*µm* × 5.5*µm*) by a tube lens (*f* = 125 *mm*). The detector is a superfast CCD camera (GZL-CL-22C5M-C, Point gray, USA) with a maximum frame rate of 2.3K *frames*/*sec* and have a quantum efficiency of 0.56. The images were collected and sent to the computer for further processing.

### B. Theoretical Investigation of System PSF

Beads flowing through the channel are conveniently modelled as Gaussian,

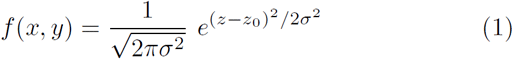

where, *z*_0_ and *σ* are respectively the mean and standard deviation.

The diameter of the bead is approximately given by,

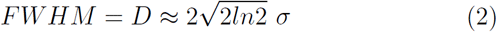

During flow, the beads undergo motion-blur at a fixed frames / sec. As a result, the beads appear elongated in the recorded image. Considering a flow-rate of *Q* through the channel of cross-section *A*, and a camera exposure of *τ*, the elongation of a point source (due to flow) is given by,

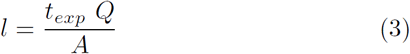

As a consequence of flow, the PSF appear elongated along the z-direction and so the changes in *σ*_*z*_ can be expressed as,

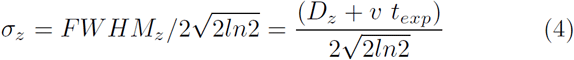

where, *D* is the diameter in static condition, and *FWHM*_*z*_ is the full-width at half-maximum of PSF along z-axis.

The resultant *σ* of the resultant Gaussian (for beads) along *y* and *z* axes are given by,

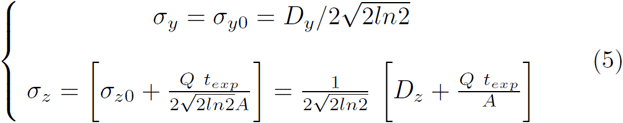

where, *v* = *Q*/*A* is the speed of fluid flowing through a channel of cross-section A at a flow-rate *Q*. Here, *σ*_*z*0_ = *σ*_*y*0_ = *σ*_0_ (say) represents standard deviation along *z* and *y* axis respectively at zero velocity (*v* = 0).

The resultant recorded bivariate Gaussian PSF can thus be expressed as,

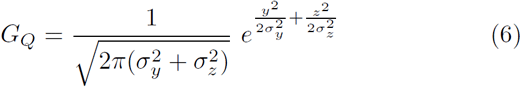

where, *σ*_*y*_ and *σ*_*z*_ are as given by eqn. 5.

Experimentally, the images of the bead sample (YZ-plane) are recorded. Assuming the beads as Gaussian, a bivariate-Gaussian function is fit to calculate the flow-induced shifts (along *z*-axis) with respect to static beads (assumed as isotropic Gaussian). Using the *σ*_*x*_ and *σ*_*y*_ parameters, a new 2D Gaussian is generated which is used as the system PSF. The process is carried out for all flow-rates. The apparent size change in system PSF along the flow direction (z-axis) at different flow-rates is detailed in Fig. 3.

The flow-variant PSFs are thus constructed from the data at every flow-rate at which the beads were flown. This PSF is then used in the deconvolution process to retrieve the object (cells) as described in the section IV C. For our case, we have taken fluorescent bead of size 1*µm* for which the emission occurs at λ_*em*_ = 575 *nm*. We have used a detection objective of *NA* = 0.4 (20X). The flow-rates vary from 50 *nl* min to 2000 *nl*/*min*.

### C. Image Reconstruction using Flow-Variant PSF

In imaging cytometry, the recorded image *g* of the object (cell) *o* flowing through the channel can be modelled as,

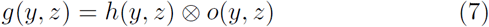

where, *h* is the PSF of the dynamic flow system.

We seek to determine the object function *o* from the recorded image. In the Fourier domain, the above equation can be expressed as,

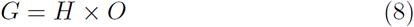

where, *G, H, O* are respectively the Fourier transform of *g, h, o*.

The object function can be retrieved from the above equation. Computationally, this is achieved by inverting the function in Fourier domain,

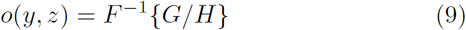

The above problem is inverse in nature and commonly known as, deconvolution.

We have used HeLa cells (of size ∼ 15 − 25 *µm*) and the mitochondria is labelled using mitotracker orange dye following the process described in section III F. The cells were flown at varying flow-rates and the images were recorded at video-rate (38 *Hz*). The ergonomic design of our system allowed collection of 4 − 8 sectional images as the cells pass through the light sheet. The 2D images were deconvolved using shift-variant PSF and images were reconstructed. Subsequently, 2D sectional images were stacked together to reconstruct the cell volume (see, Fig. 6).

### D. Microfluidic Chip Fabrication

Using Clewin 4, channel features are designed on 4-inch silicon disc. Negative mask is printed from Clewin 4.gbr file. Master mold fabrication is done in clean room facilities at Nanoscience Facility, Indian Institute of Science, Bangalore, India. Subsequently, Y-type microfluidic chips were fabricated using standard protocol. Silicon elastomer and curing reagent are mixed thoroughly in the ratio of 10:1. A net mixture amount of 33gm is dicicated with vacuum pump for 20 minutes until air bubbles are removed. Decicated mixture is gently poured on the top of master mold and is cured in hot oven at 60 degree centigrade for 3 hours. Cured PDMS is peeled off from master mold and useful region are extracted from it by cutting. Thus, replica of the micro-channels on the PDMS blocks are obtained. Inlet and outlets are punched with 1.0 mm diameter PDMS puncher and cleaned with isopropanol and acetone. Washed PDMS with microchannel channel and bonding glass (0.15 mm thickness) are plasma cleaned for 5 minutes. As soon as plasma cleaning is completed PDMS is placed on the top of coverslips. Following baked on hotplate for 5 minutes at 90-degree centigrade. Using microfluidic Teflon tubing (inner diameter of 0.5 mm) reservoirs is connected to inlet and outlet is connected to flow pump (New Era Flow Pump, Model No: NE-1002X). Microfluidic chips were ensured leakage-free by flowing distilled water while the features of micro-channels are obtained using low concentration TRITC solution (*Ex*/*Em* = 557/576*nm*) before actual imaging with beads and HeLa cells.

### E. Flow pump and Data acquisition

The 2D sectional images of HeLa cells were recorded by the CCD camera. Two major system were synchronized to record the data, (1) flow control system and (2) Lab-view Image Acquisition. The flow is controlled using a flow-pump operated in a withdrawl mode, and at flow-rates ranging from 500 − 2000 *nl*/*min* (for HeLa cells).

Needle of flow-pump (BD,1ml conventional syringe of diameter 4.80mm) is attached to outlet tube of the microfluidic device to withdraw the cell embedded solution from the reservoir through the micro-channels. Volumetric flow rate (in nano litter cube) is the set point on pump. Dispense module is activated for withdrawal-mode of the pump. The pump was operated from 500 *nl*/*min* to about 2000*nl*/*min* for data acquisition.

Raw image data are recorded using a CCD camera (Point gray camera, Model:GZL-CL-22C5M-C, Point gray, USA). For accurate and fast data collection the camera is interface with NI PCI 1433 card (frame-graber device) and is programmed using LabVIEW-12. Set point on the program is exposure time and camera resolution. In terms lightsheet thickness, optimum value of exposure time as decided by the thickness of light sheet and flow-rate (see, eqn.(3)) and camera characteristics (full window size: 2048 × 1088, pixel size = 5.5*µm*) are set for imaging. Raw Data recorded are saved in.bmp format.

### F. Sample Preparation

#### 1. Fluorescent Beads

Beads were used for both calibration and as a test sample. 1*µl* of Invitrogen FluoSpheres Carboxylate-Modified Microspheres, 1.0*µm* diameter (Nile Red fluorescent, *Ex*: 535 *nm*/*Em*: 575 *nm*) (Invitrogen, USA)is suspended in 1 *ml* of distilled water. Through mixing is done by pipetting and the mixture is loaded to the sample reservoir of parallel iLIFE system for counting, imaging and parameter estimation.

#### 2. Cell line and maintenance

HeLa cells (human cervical carcinoma cell line) were used for the experiment. It was grown as monolayers in Dulbecco’s modified minimal Eagle’s medium (DMEM) (Gibco, Thermo Fisher Scien-tific) supplemented with 10% FBS(Gibco, Thermo Fisher Scientific) and 1% penicillin –streptomycin solution (Gibco, Thermo Fisher Scientific) and incubated in humidified chamber at 37°*C* and 5%*CO*2 (Thermofisher, Forma series 381). The cells were thawn and standard running culture is maintained. After 2 passage, the cells were prepared for experiment. Hemocytometer is used to count cells after every passage and approximately 100,000 cells were maintained. The cells were passaged in every 2-3 days to maintain healthy cell-lines.

#### 3. Mitochondrial labelling using mitotracker orange

Cells were seeded (10^5^ cells) in T25 flasks 24 h before mitochondrial labelling. Mitotracker orange (purchased from Invitrogen) of 1mM concentration in DMSO was used as stock. Subsequently, 6 different working concentration of mitotracker orange (100,125,150,175,200,225 nM) in DMEM was tested to find the optimum concentration with respect to the density of HeLa cell, staining efficiency and quantum yield in imaging flow cytometry. Cell with 70% confluency was taken and washed with PBS to remove cell debris. It was incubated with Mito tracker orange of 175nM concentration and kept for 1h 15 min in the incubator with 5% CO2 and 37°*C*. After incubation, cells were trypsinized to detach them from disc-surface and the cells were centrifuged at 2000 RPM for 5min. Subsequently, the cells were resuspended in PBS and prepared for experiments. The cells were loaded in the sample reservoir and flown through microfluidic chip for volume imaging cytometry.

## V. ACKNOWLEDGMENT

We acknowledge financial support from Department of Biotechnology, New Delhi, India.

## Author Contributions

PK and PPM have conceived the idea and designed the experiment. PK, PJ and PPM have performed the experiments. PJ and JB has prepared the bead sample and fluorescently-labelled HeLa cells. PK, PJ, JB and PPM have carried out the analysis. PPM, PK and PJ have written the manuscript.

## Additional Information

The author declare no competing financial interest.

